## Supplementary figure 1-18 for "Simultaneous cyclin D1 overexpression and p27^kip1^ knockdown enable robust Müller glia cell cycle reactivation in uninjured mouse retina"

#### Supplementary Fig. 1

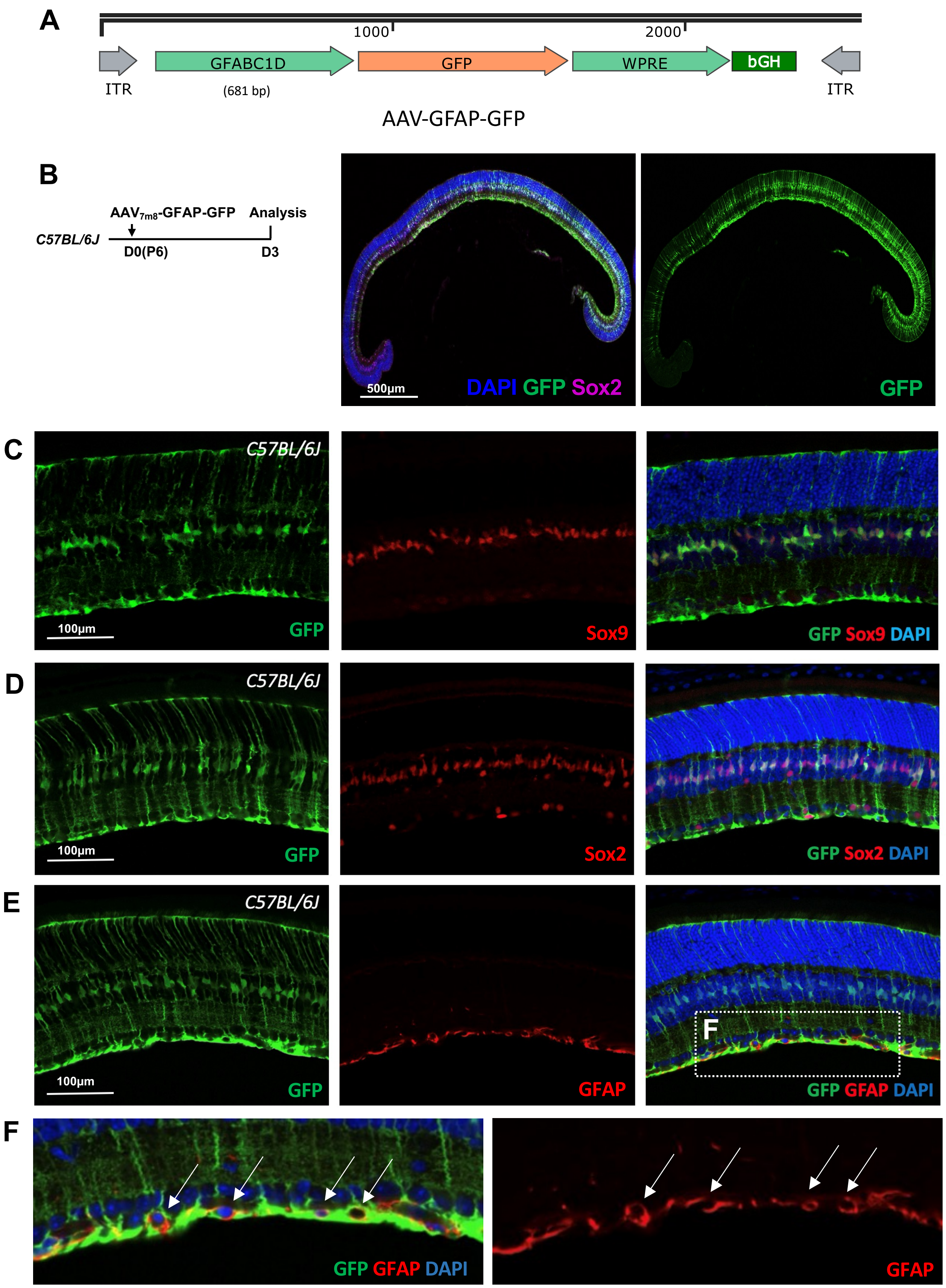

#### Supplementary Fig. 2

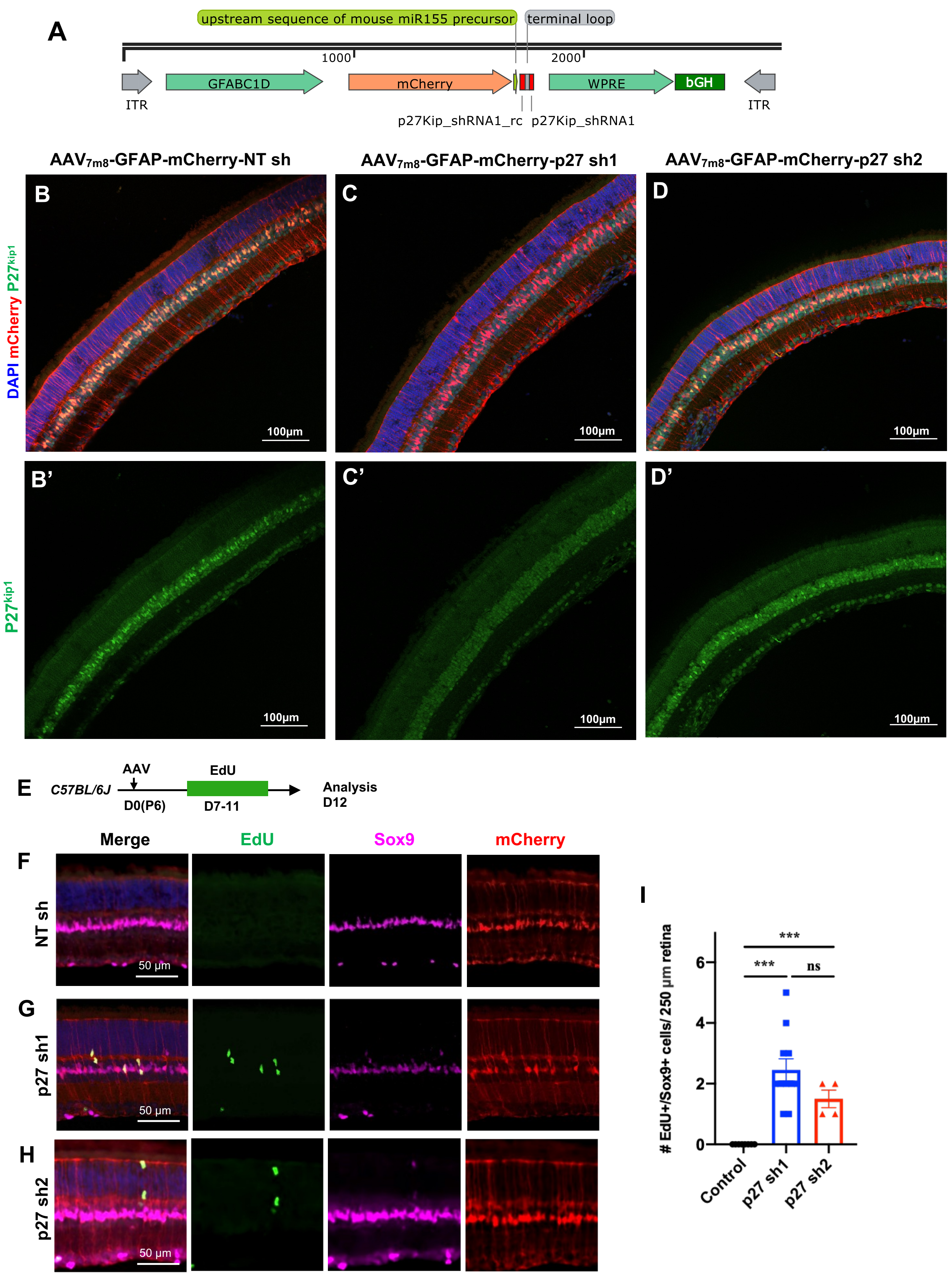

#### Supplementary Fig. 3

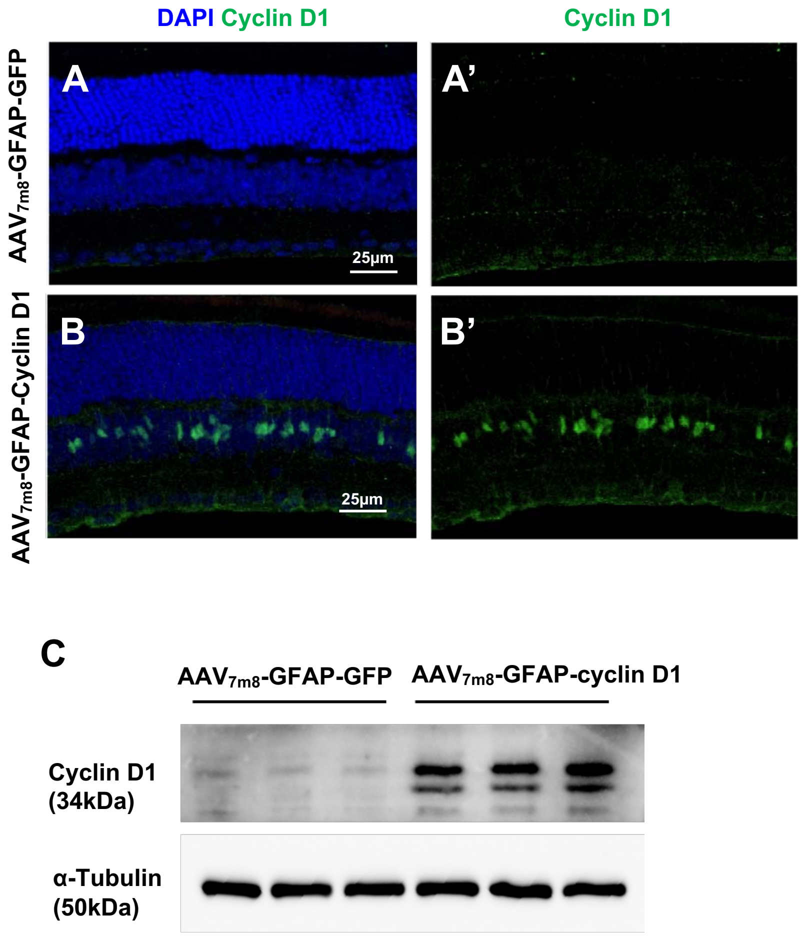

**Supplementary Fig. 4**

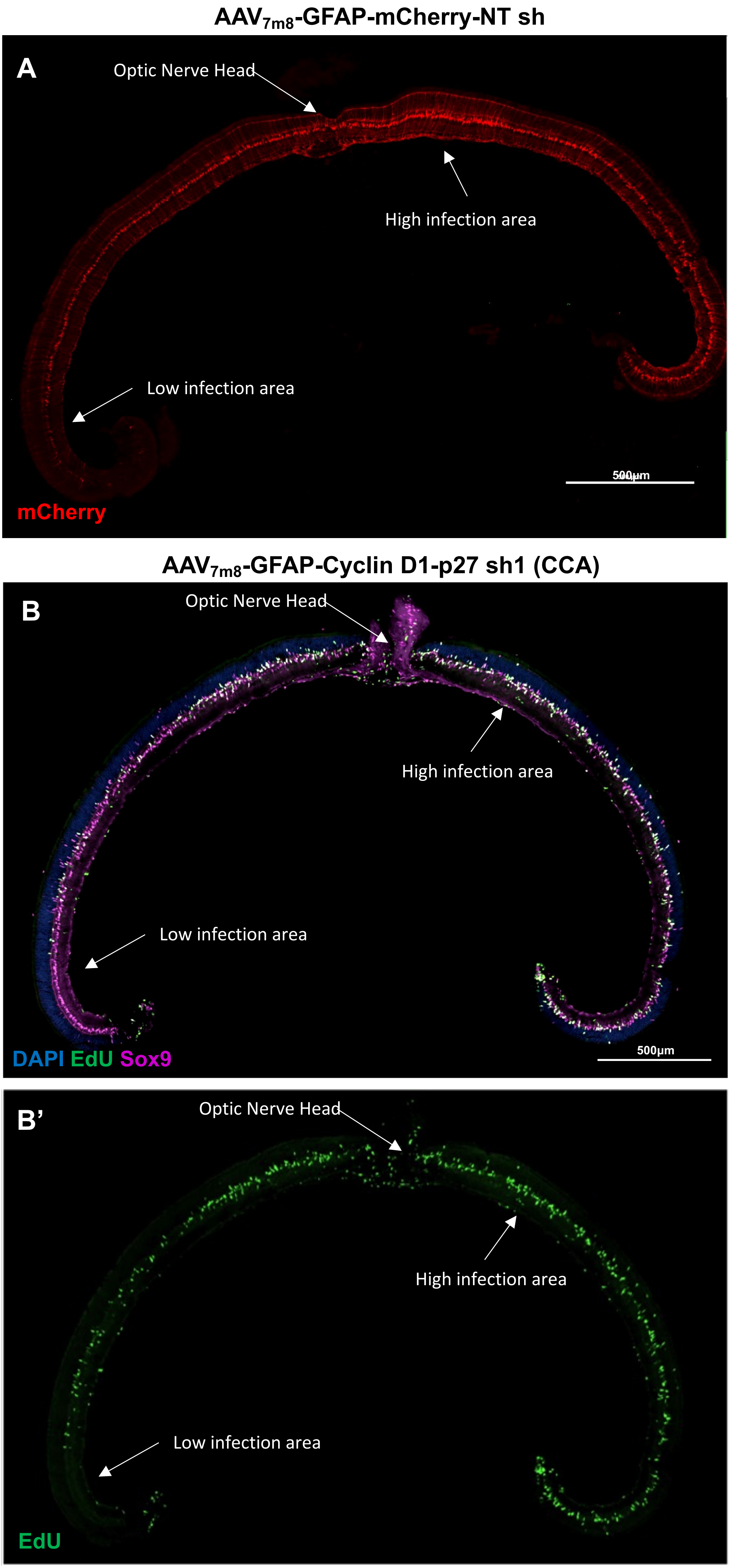

#### Supplementary Fig. 5

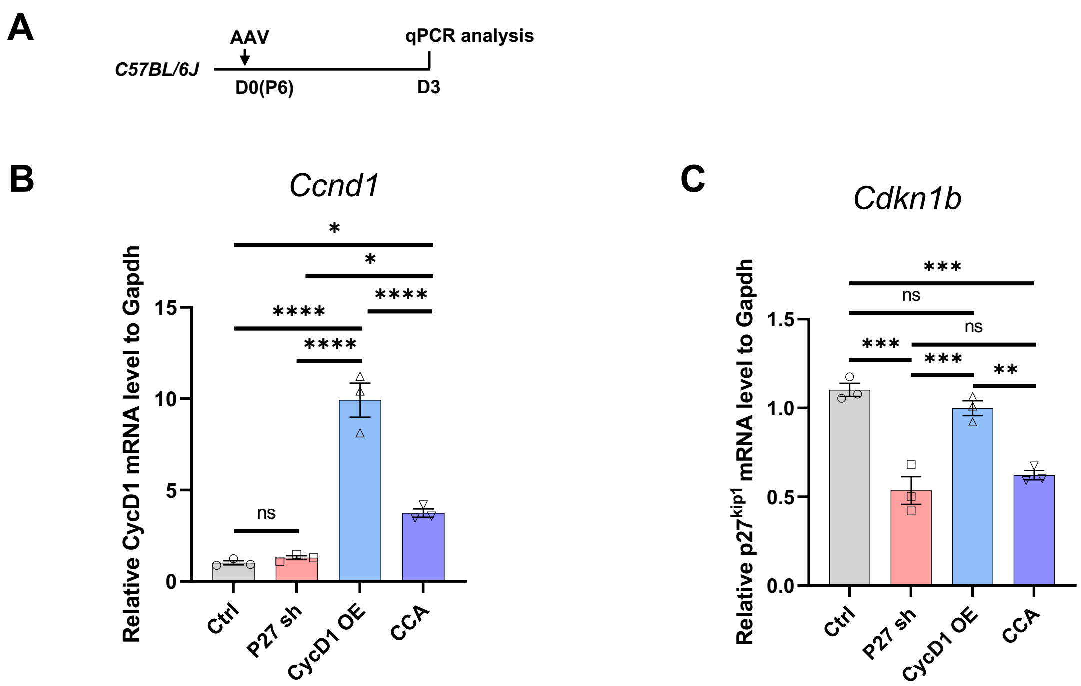

#### Supplementary Fig. 6

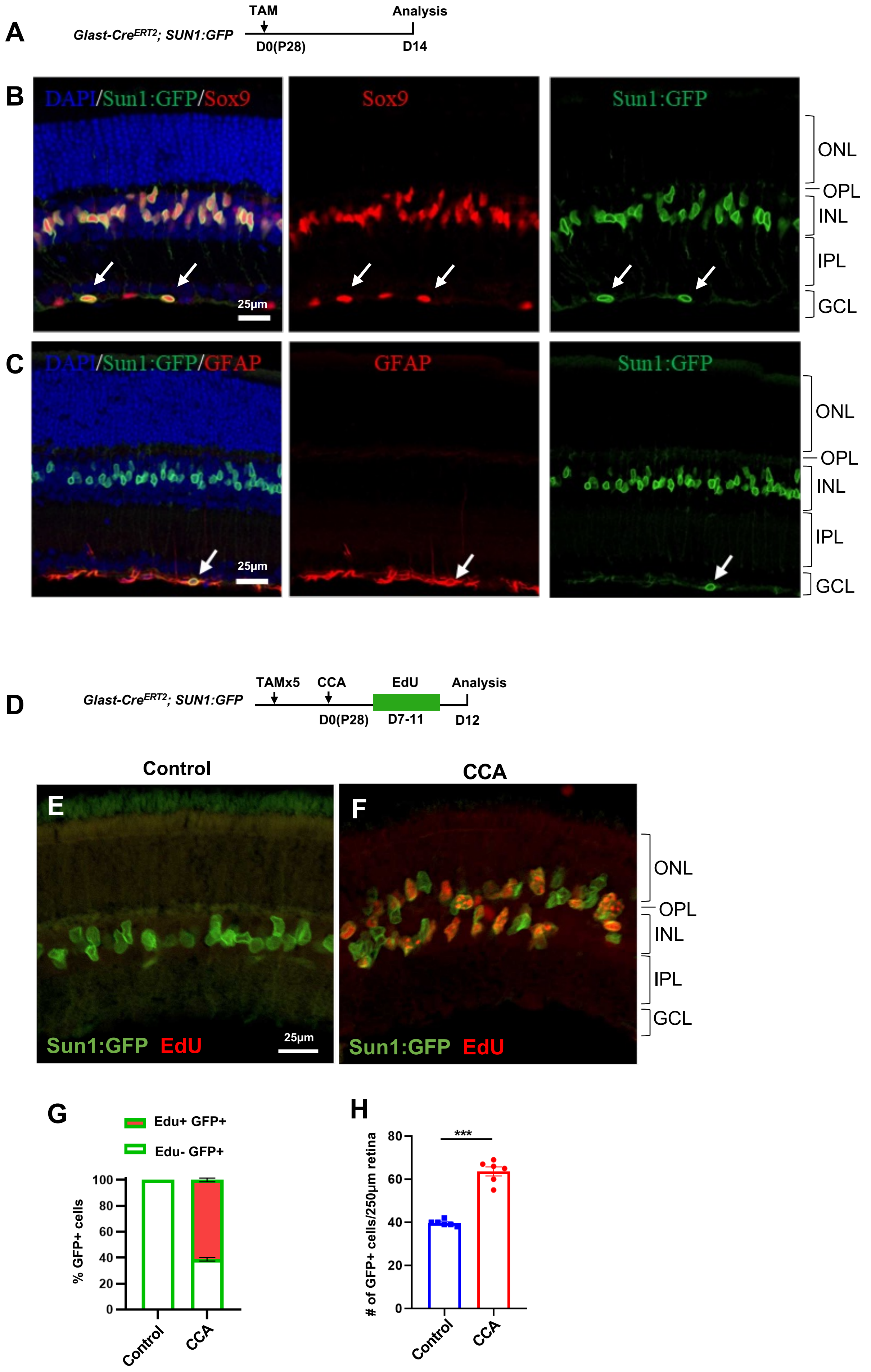

#### Supplementary Fig. 7

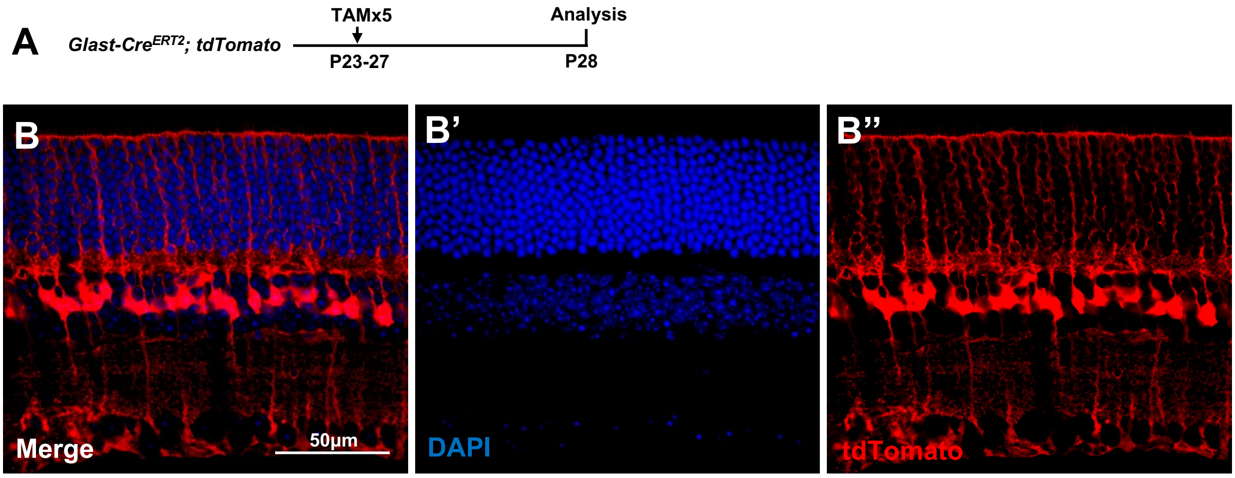


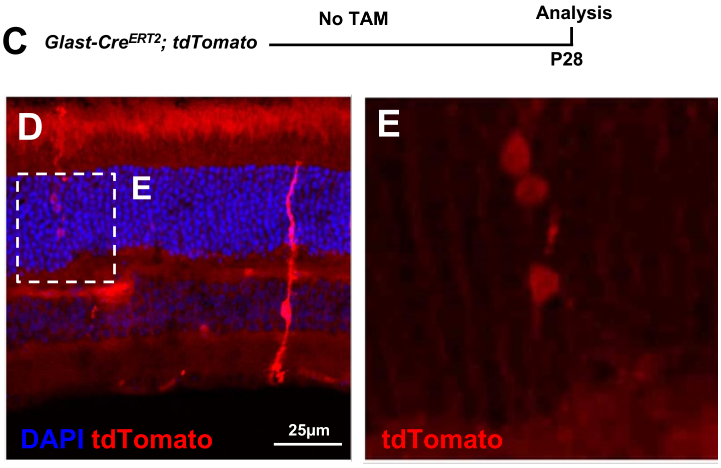


**F**

| | # of cells with leaky tdTomato expression * |
|---|---|
| MG | 34 |
| rod | 14 |
| Cone | 0 |
| BP | 0 |
| AC | 0 |
| HC | 0 |
| RGC | 0 |

* Screened 8 retinas (total of 128 whole retinal cross-sections)

**Supplementary Fig. 8**

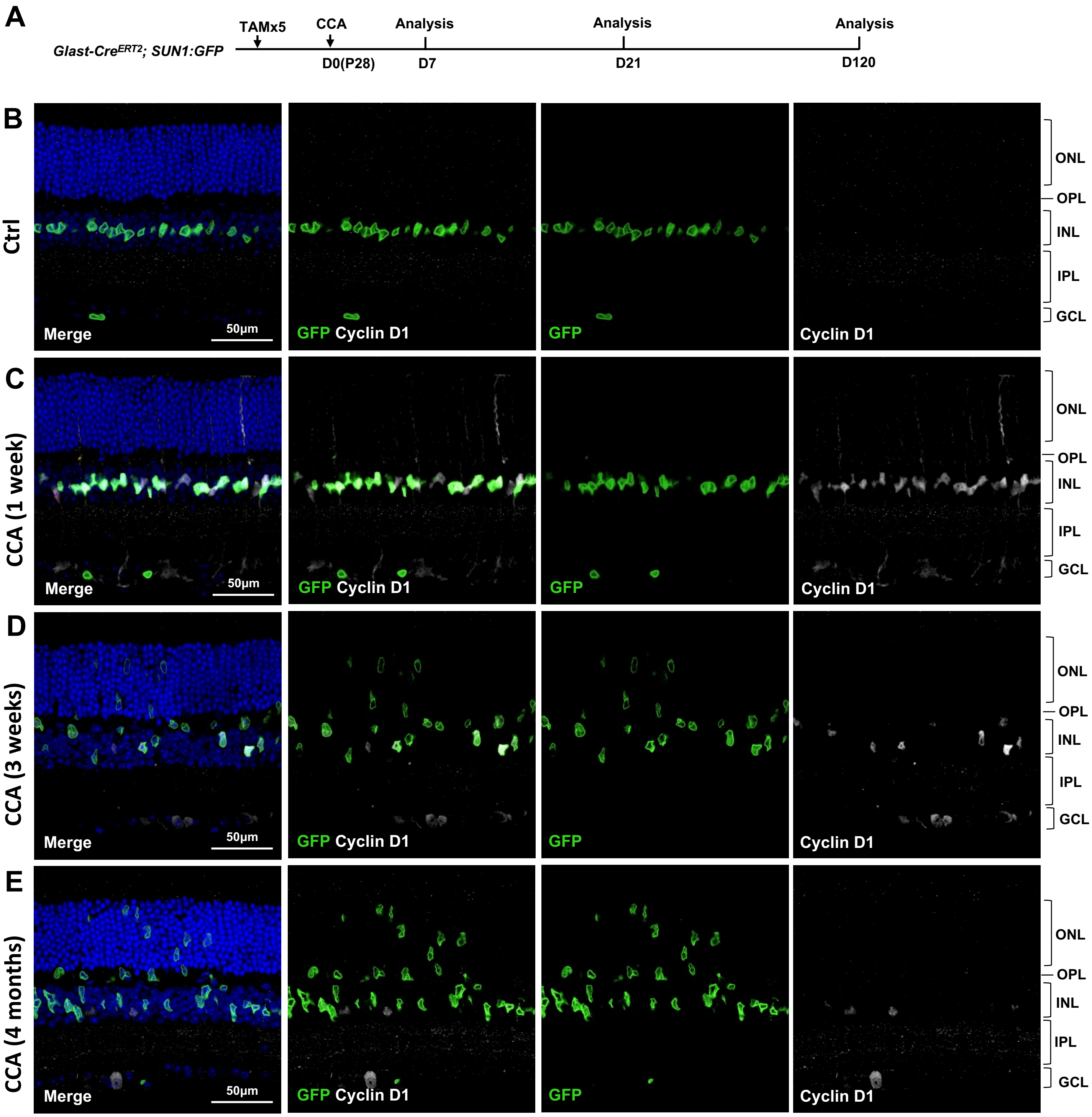

#### Supplementary Fig. 9

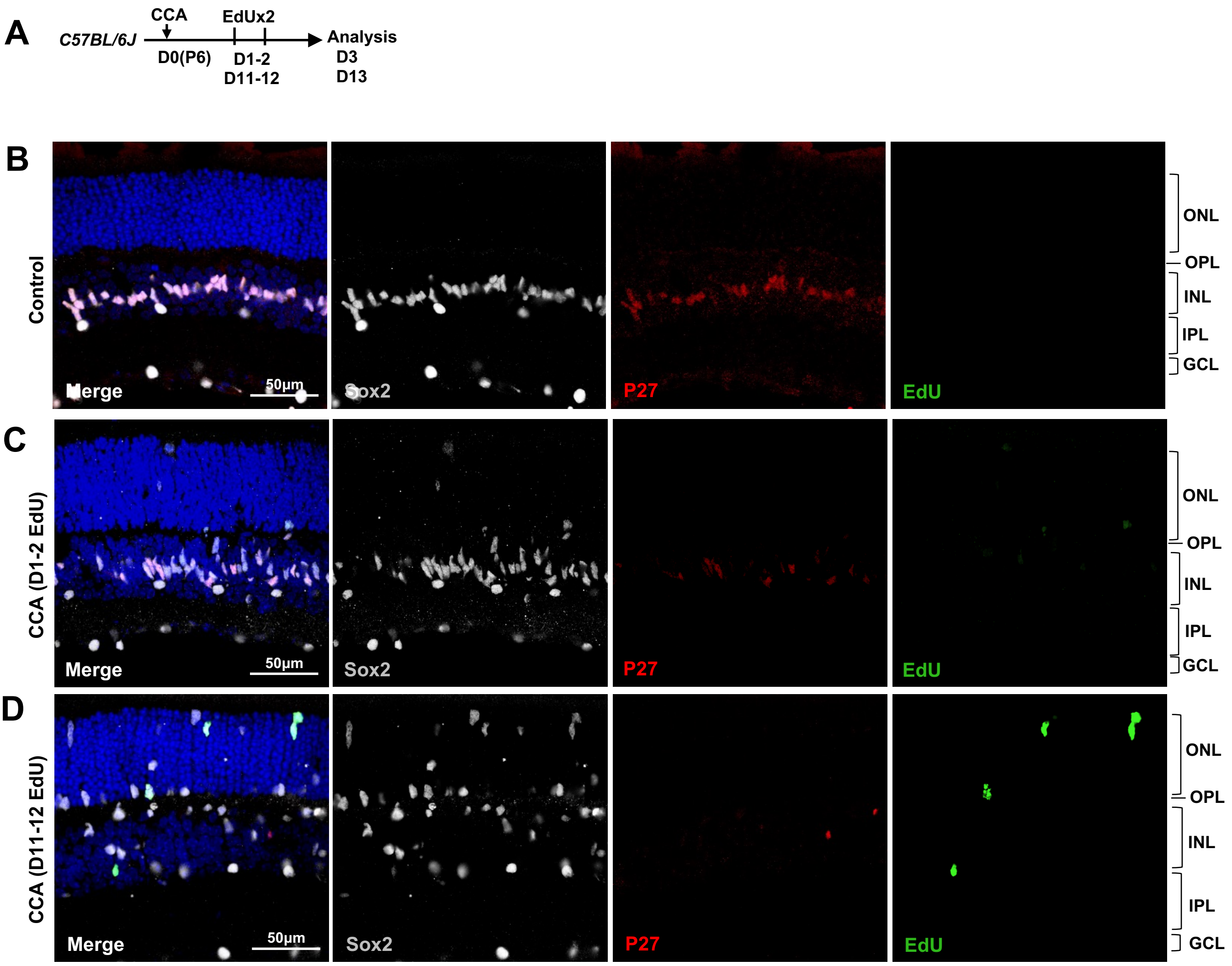

#### Supplementary Fig. 10

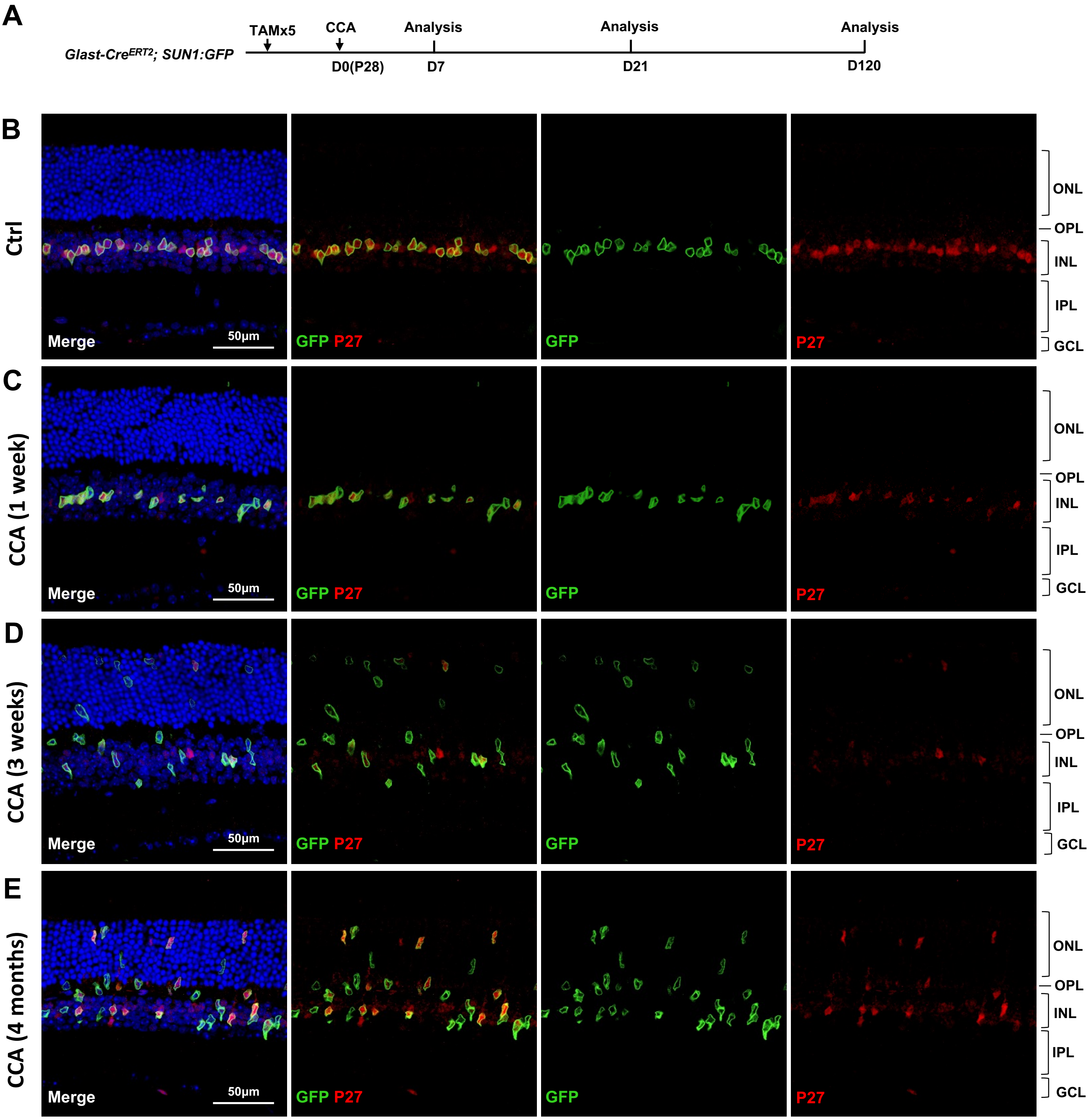

**Supplementary Fig. 11**

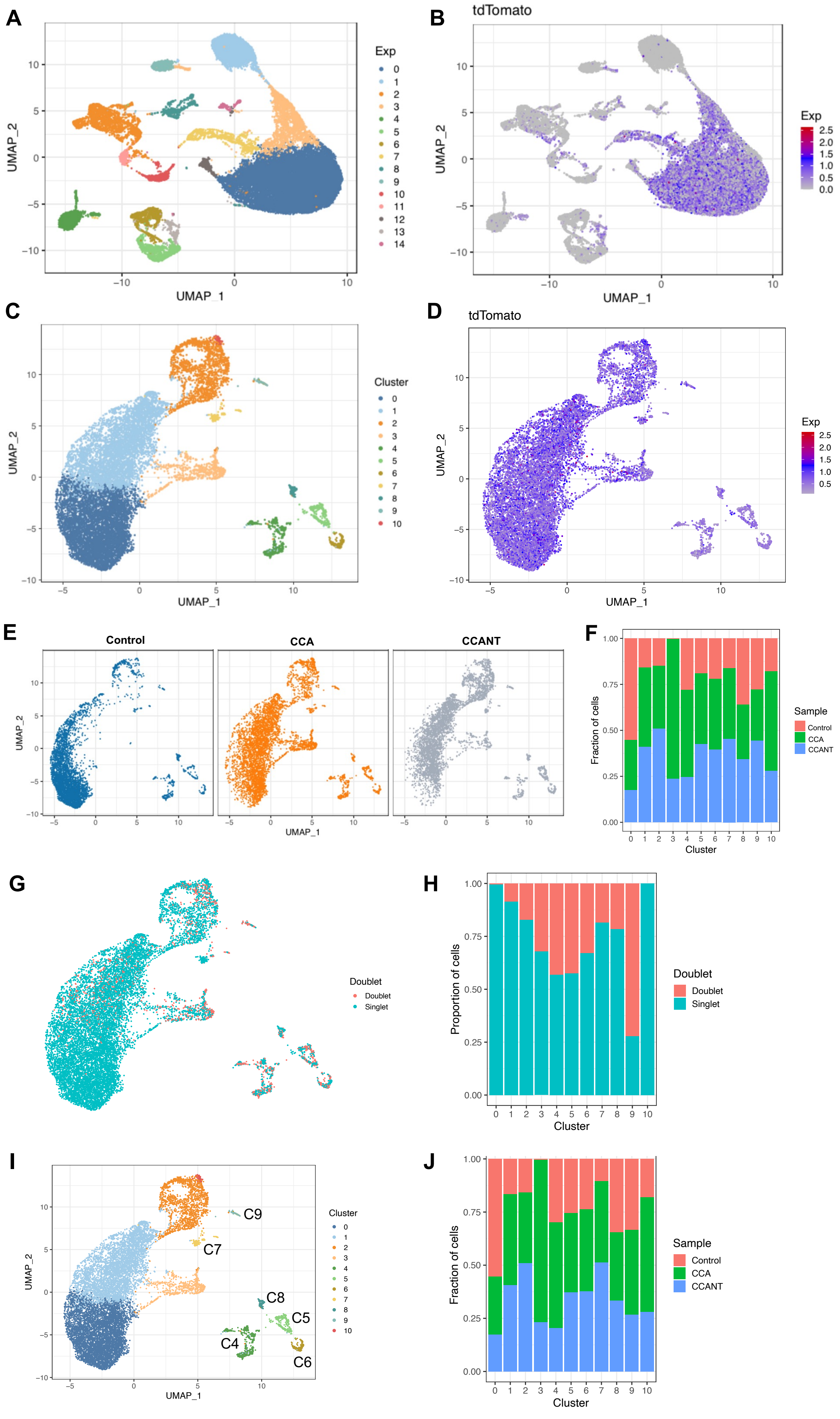

### Supplementary Fig. 12

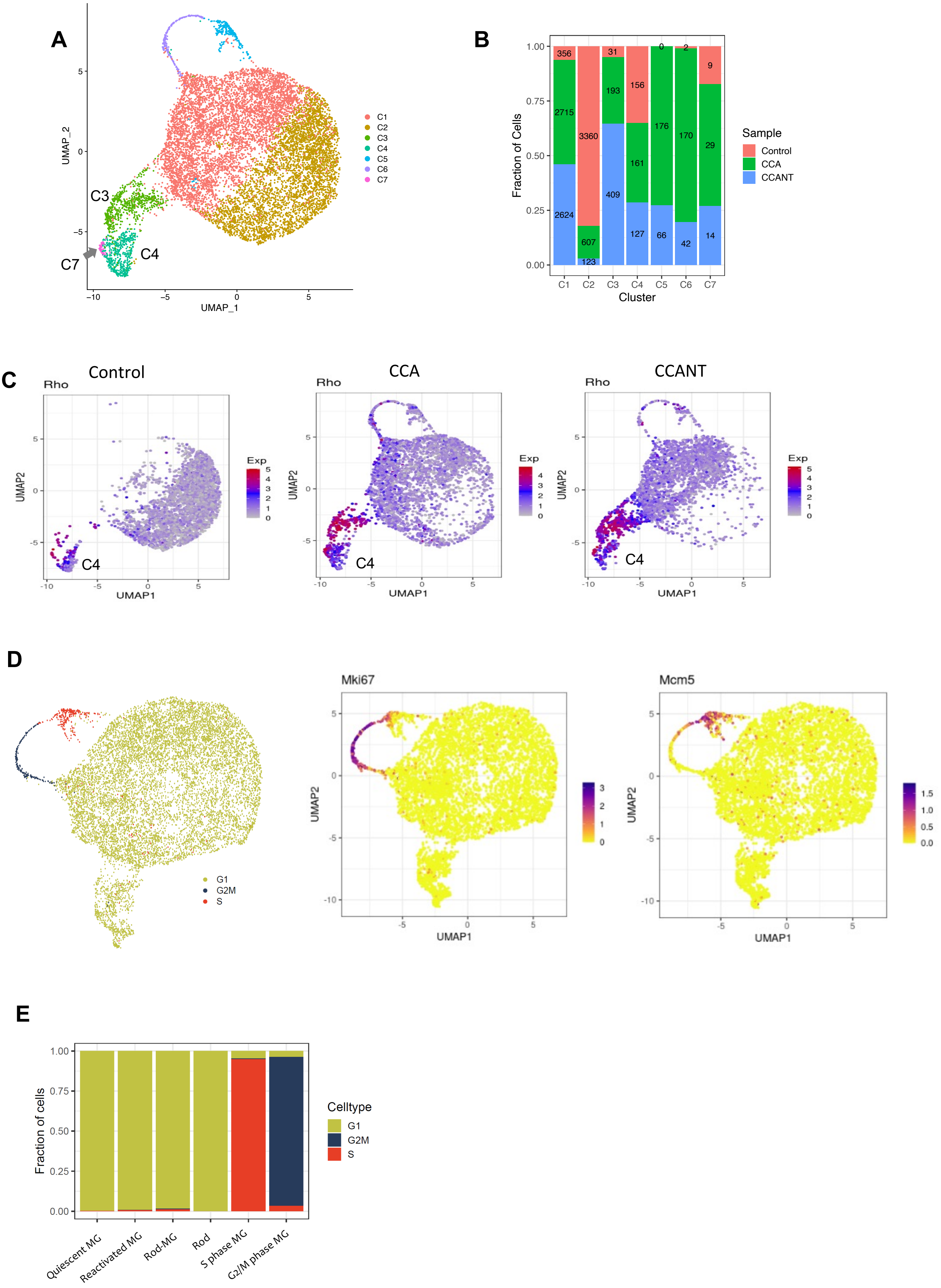

**Supplementary Fig. 13**

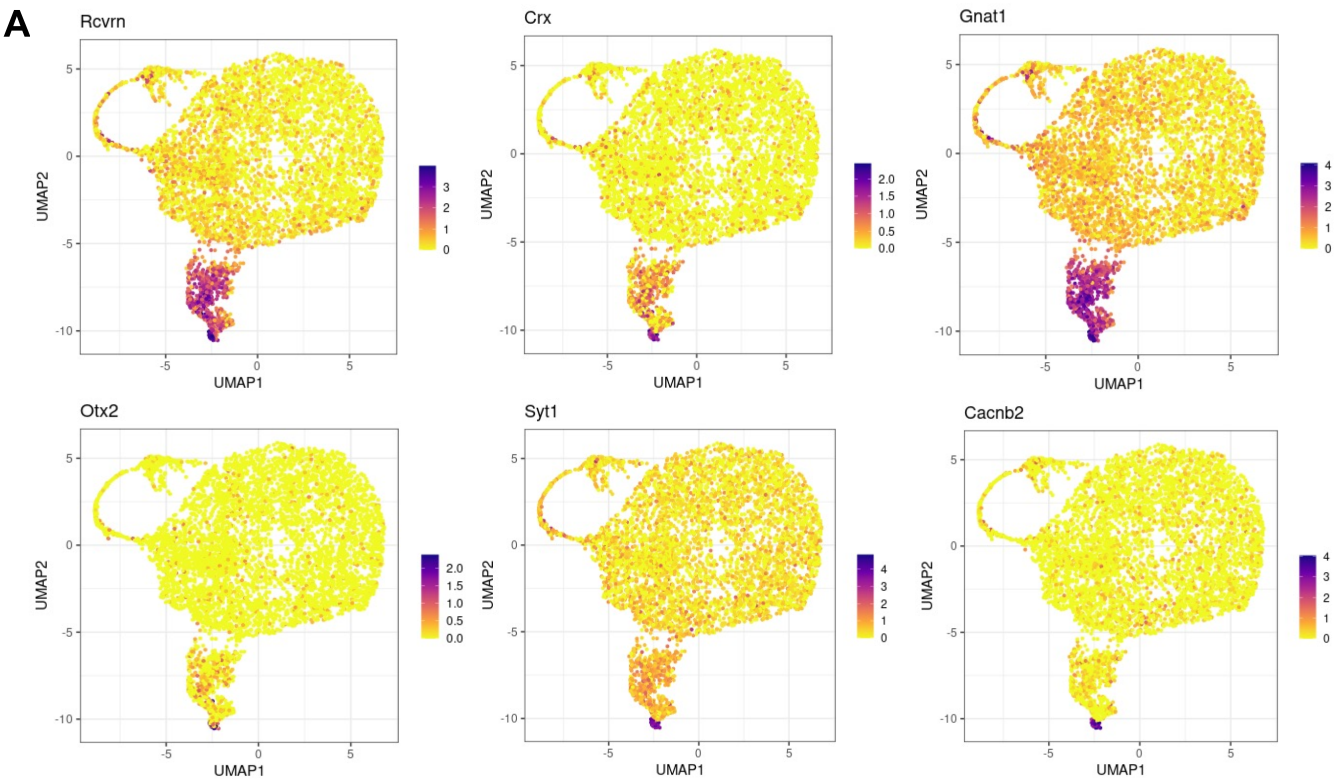

#### Supplementary Fig. 14

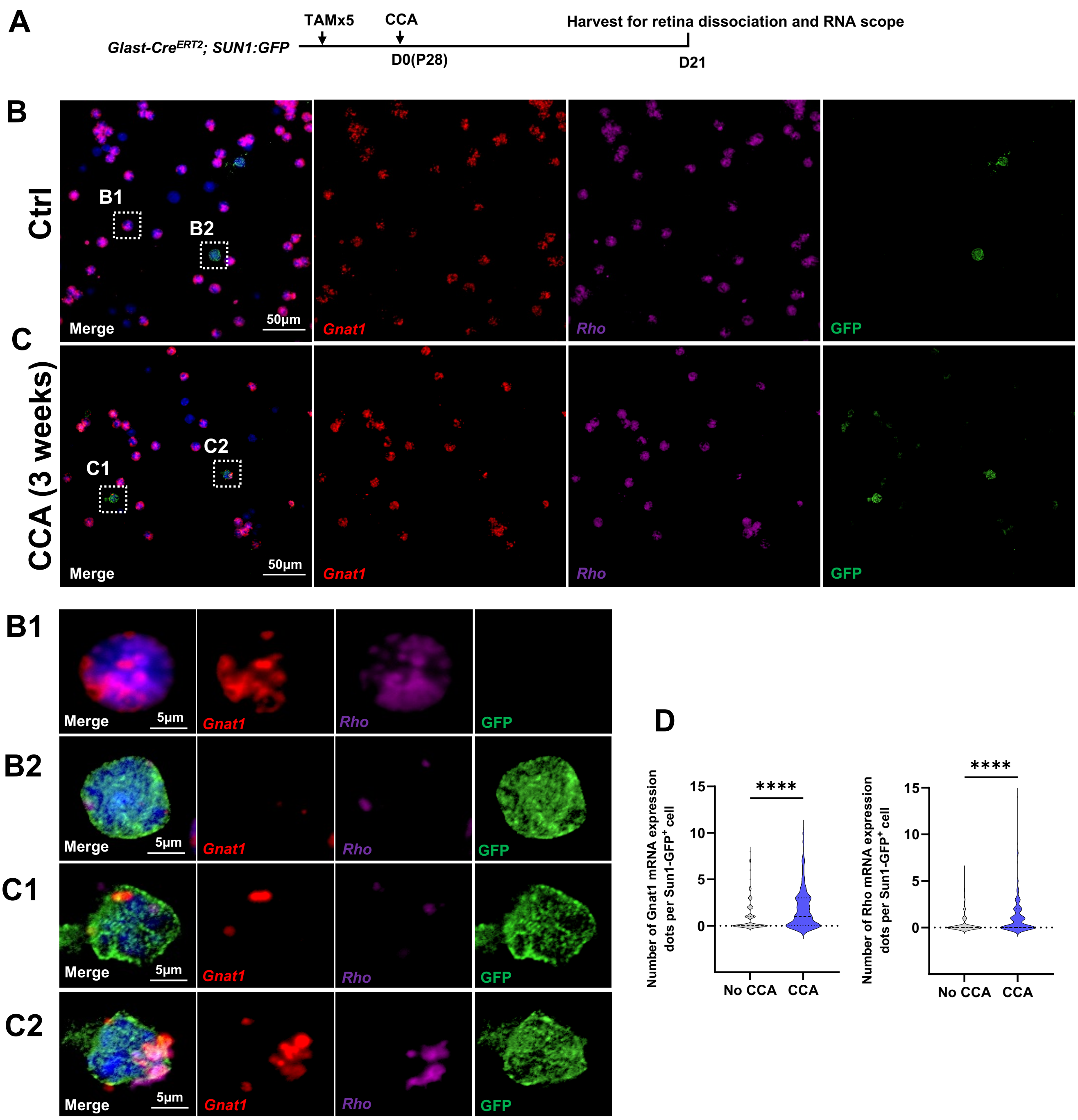

### Supplementary Fig. 15

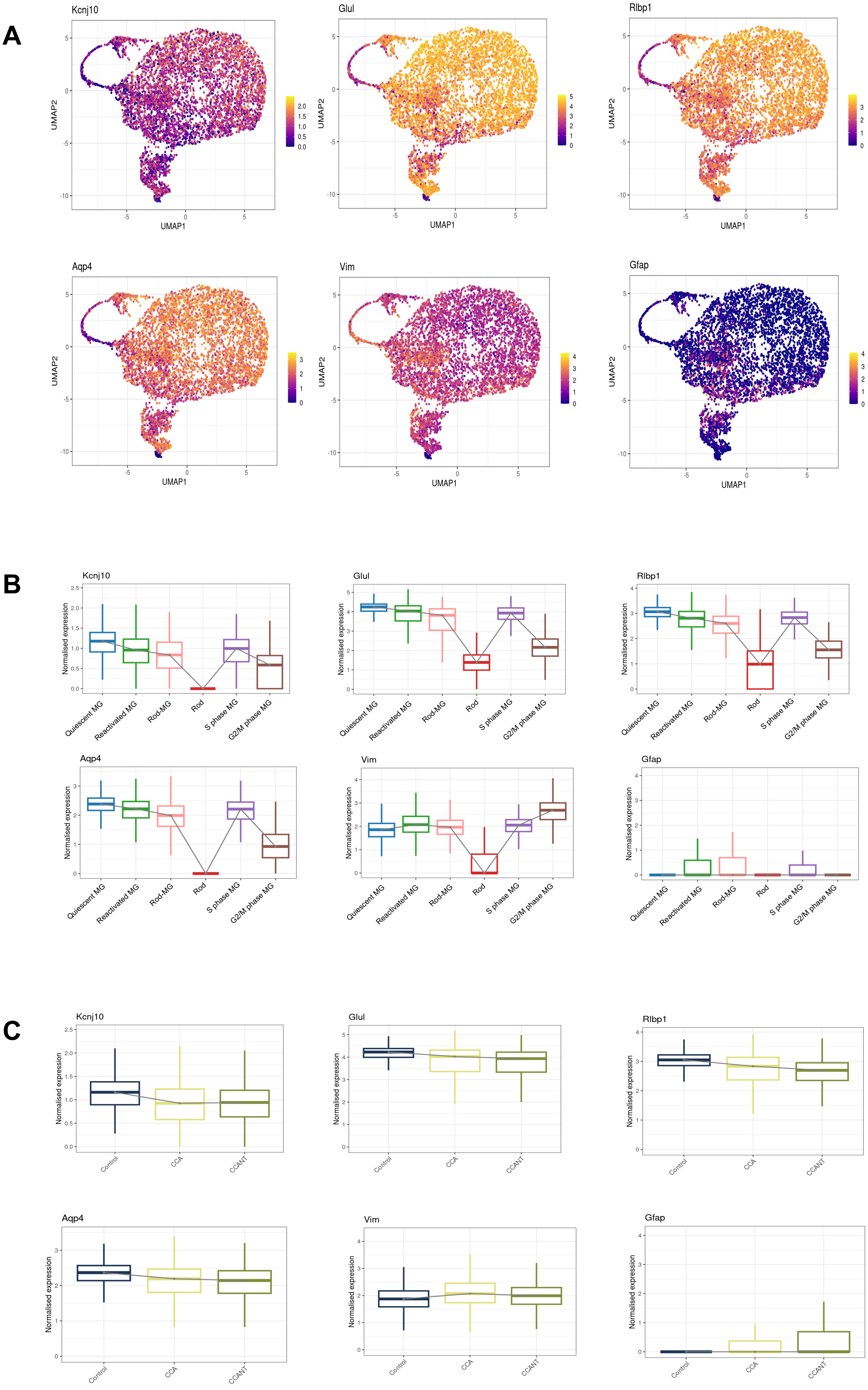

#### Supplementary Fig. 16

**A**

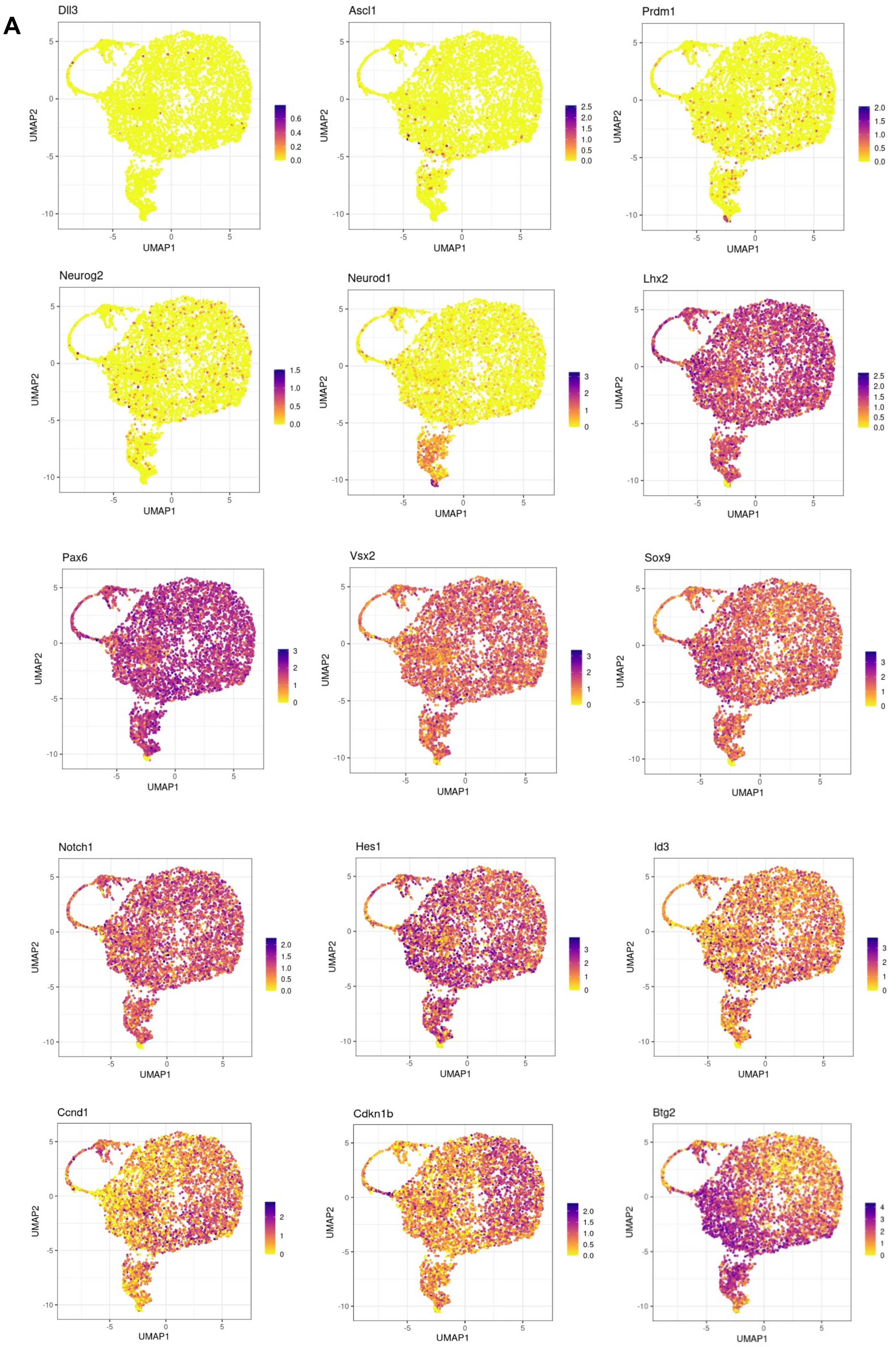

### Supplementary Fig. 17

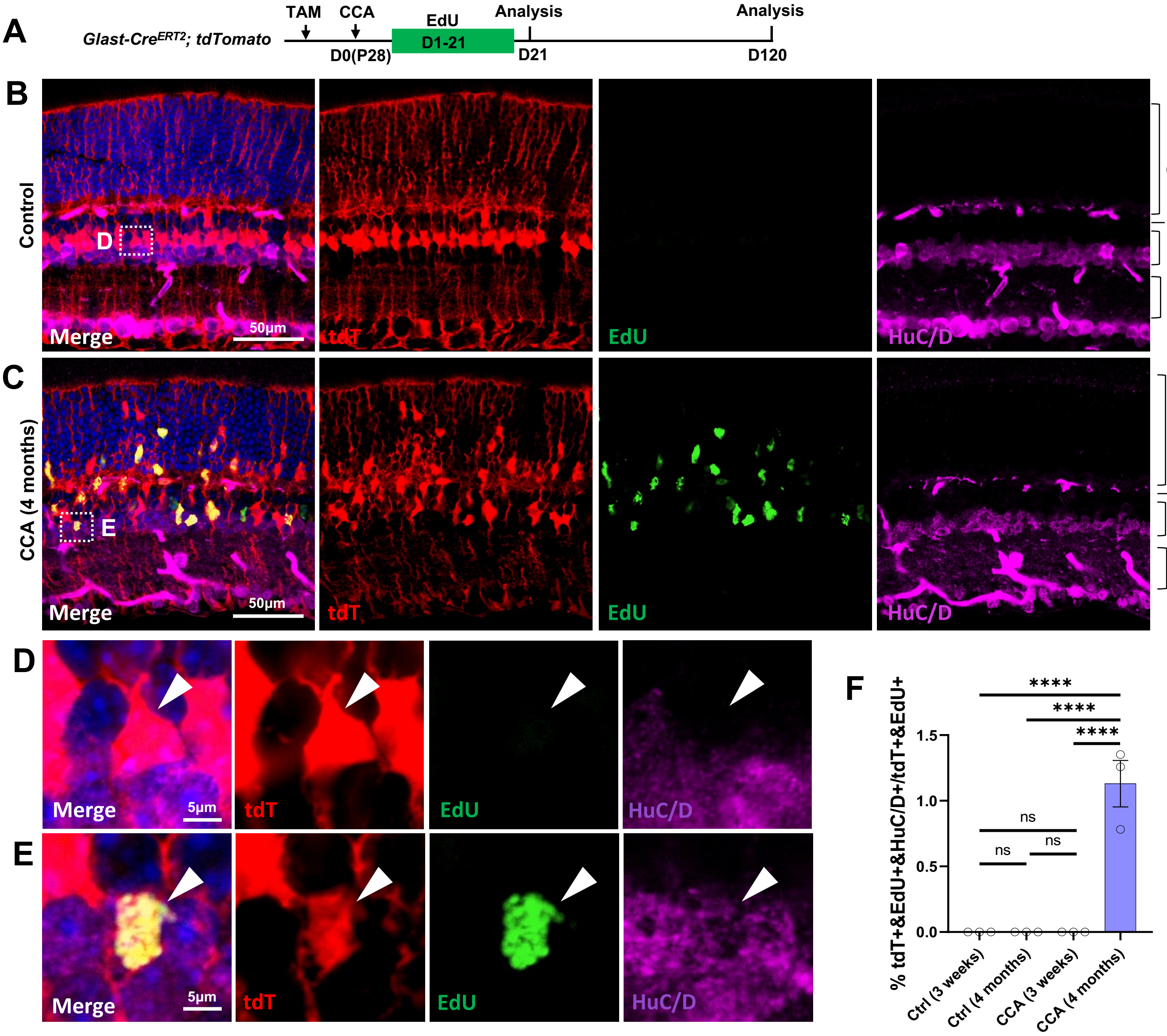

### Supplementary Fig. 18

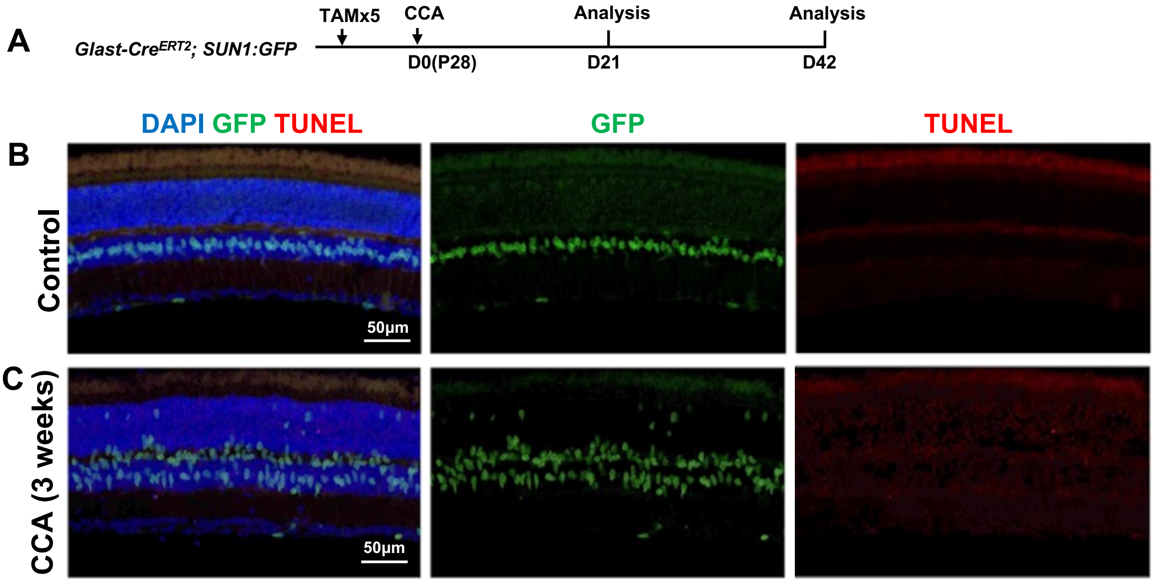
