## Supplementary Information for "Simultaneous cyclin D1 overexpression and p27^kip1^ knockdown enable robust Müller glia cell cycle reactivation in uninjured mouse retina"

**Supplementary Methods**

**Quantitative PCR**

RNA was extracted from whole retina using Trizol (Thermo Fisher Scientific) followed by the Quick-RNA MicroPrep Kit (Zymo Research). RNAs were converted to cDNA using a PrimeScript RT reagent kit with gDNA Eraser (Takara Bio). qPCR was performed using the PowerUp Sybr Green Master Mix (Thermo Fisher Scientific) on QuantStudio 3 Real-Time

PCR stems (Applied Biosystems). Gapdh was used as the normalizingcontrol. qPCR primers are listed below:

p27_qPCR_F: TCAAACGTGAGAGTGTCTAACG

p27_qPCR_R: CCGGGCCGAAGAGATTTCTG

CycD1_qPCR_F: CCCAACAACTTCCTCTCCTG

CycD1_qPCR_R: TCCAGAAGGGCTTCAATCTG

Gapdh_qPCR_F: AGGTCGGTGTGAACGGATTTG

Gapdh_qPCR_R: TGTAGACCATGTAGTTGAGGTCA

**RNA *in situ* hybridization on dissociated retinal cells**

Fresh retina was dissociated in Papain (Worthington) and gently pipetting using a 1 ml pipette tip, and the dissociated retina cells were then filtered with 40 µm strainer (pluriStrainer). Filtered retinal cells were then seeded and cultured in a chamber slide (Thermofisher) at 37°C in a cell incubator. Following incubation, the cultured retinal cells were washed with PBS and fixed with 4% PFA. Dissociated cells were stained with GFP antibody (AB_2307313; Aves Labs) at 4 °C overnight and then hybridized with RNA probes (Advanced Cell Diagnostics Cat.No. 474801-C3 for Mm *Rho*, 524881-C2 for Mm *Gnat1*) for 2 hours at 40 °C for two hours. Following *in situ* RNA hybridization steps, slides were stained using secondary antibodies (Jackson ImmunoResearch) and DAPI for two hours at room temperature. The fluorescent signals were visualized and captured using Nikon A1HD25 High speed and Large Field of View Confocal Microscope. The mRNA levels were quantified by counting the numbers of RNA dots using ImageJ by another experimenter with group masked.

**Supplementary Table 1**

Differential gene expression analysis between different cell types or three treatment groups from the scRNA-seq data.

**Supplementary Figure legends**

**Supplementary Figure 1. Specific GFP expression in MG of mouse eyes intravitreally injected with AAV-GFAP-GFP vector.**

(A) Schematic diagram of the AAV-GFAP-GFP vector genome. The promoter sequence was the GFABC_1_D region of human glial fibrillary acidic protein (GFAP) gene promoter. (B) Mouse retinas infected by AAV-GFAP-GFP virus via intravitreally injection on P6, and GFP expression started as early as three days post-injection. (C-F) Immunohistochemistry of MG marker Sox9 (C) and Sox2 (D), as well as astrocyte marker GFAP (E-F). (F) Zoomed-in image of the boxed area in E’’, illustrating that GFP+ MG end feet encircle GFAP-stained astrocytes (arrows). GFP signal and the GFAP staining did not colocalize, indicating that AAV-GFAP-GFP specifically label MG rather than astrocytes.

**Supplementary Figure 2. Downregulation of p27^kip1^ expression in MG by AAV-GFAP-mCherry-p27 shRNA1/2 vector**

(A) Schematic diagram of the AAV-GFAP-mCherry-p27 shRNA vector genome. P27 shRNA was cloned in the mouse miR155 cassette and inserted in the 3’UTR of mCherry. (B-D) Immunohistochemistry for p27^kip1^ in retinas infected with AAV-GFAP-mCherry-NT sh (B), AAV-GFAP-mCherry-p27 sh1 (C), and AAV-GFAP-mCherry-p27 sh2 (D). p27 sh1 with higher p27^kip1^ knockdown efficiency was used in this study. (E) Experimental design of EdU assay. (F-H) Representative images of retinas infected with various AAV vectors expressing different shRNAs. (J) Quantification of EdU+ Sox9+ cells in retinas injected with AAV-NT shRNA (n=8), AAV-p27 shRNA1 (n=11), and AAV-p27 shRNA2 (n=4). Data presented as mean ± SEM. ns=not significant, ****P*<0.001 (one-way ANOVA analysis with post-hoc Tukey test).

**Supplementary Figure** **3. MG-specific upregulation of cyclin D1 in mouse retina infected with AAV-GFAP-cyclin D1**

(A-B) Immunostaining for cyclin D1 in retinas infected with either control AAV-GFAP-GFP vector (A) or AAV-GFAP-cyclin D1 (B). (C) Western blot analysis confirming cyclin D1 overexpression in AAV-GFAP-cyclin D1-injected retinas. The retinal lysate from one mouse retina per lane.

**Supplementary Figure 4. Infection pattern of intravitreally delivered AAV vector.**

(A) Whole retinal cross-section demonstrating typical infection pattern by AAV-GFAP-mCherry-NT shRNA vector, with strong mCherry expression indicating high infection efficiency in the central retina. (B) Distribution of EdU+ cells across the retinal section, showing non-uniformity. Mouse eyes were infected by CCA vector via intravitreal injection on P6, and proliferating MG were labeled by five EdU injections from day 7 to 11 post-CCA injection. EdU+ cell hotspot typically located near the optic nerve head.

**Supplementary Figure** **5. Comparison of cyclin D1 overexpression and p27^kip1^ knockdown efficiencies by different viruses.**

(A) Experimental design. (B-C) Quantitative PCR results for *Ccnd1* (B) and *Cdkn1b* (C) in retinas from different treatment groups, showing more efficient cyclin D1 overexpression driven by AAV-GFAP-cyclin D1 compared to CCA. p27^kip1^ knockdown efficiency by AAV-GFAP-mCherry-p27 shRNA1 was comparable to that of CCA. Data presented as mean ± SEM, n=3. ns=not significant, **P*<0.05，***P*<0.01, ****P*<0.001, *****P*<0.0001 (one-way ANOVA analysis with post-hoc Tukey test).

**Supplementary Figure** **6. Characterization of MG proliferation in *Glast-Cre^ERT2^; Sun1:GFP* transgenic mice**

(A) Experimental design. (B-C) Immunohistochemistry against MG marker Sox9 (B) and astrocyte marker GFAP (C) in retinas of *Glast-Cre^ERT2^; Sun1:GFP* mice, where GFAP expression was low in MG in uninjured retinas. Arrows indicate Sun1:GFP positive astrocytes in the ganglion cell layer (GCL), which expressed high levels of GFAP. (D) Experimental design. (E-F) EdU incorporation analysis in retinas following control (E) or CCA (F) treatment. (G) Quantification of EdU+GFP+ and EdU-GFP+ cell percentages in high infection areas of *Glast-Cre^ERT2^; Sun1:GFP* retinas injected with CCA. (H) Quantification of total GFP+ MG per 250 µm in high infection areas. Data presented as mean ± SEM, n=6. *** *P*<0.001 (unpaired two-tailed student’s t-test).

**Supplementary Figure** **7. Characterization of MG labeling and leaky expression in *Glast-Cre^ERT2^; tdTomato* transgenic mice**

(A) (A) Experimental design. (B) MG were labeled by tdTomato in retinas of *Glast-Cre^ERT2^; tdTomato* mice treated with five tamoxifen injections from P23 to P27. (C) Experimental design. (D-E) Characterization of leaky expression of *Glast-Cre^ERT2^;tdTomato* mouse retina without tamoxifen induction. The boxed area in B is enlarged in C. Leaky expression was observed in MG as well as rods at very low incidence. (D) Quantification of tdTomato-positive cells in retina without TAM injection. A total of 128 retinal sections from 8 retinas were examined. 34 MGs and 14 rod cells labelled by leaky tdTomato expression were found in all sections after careful examination.

**Supplementary Figure** **8. Analysis of cyclin D1 expression at different days post-CCA injection on P28**

(A) Experimental design. (B-E) Immunohistochemical analysis of cyclin D1 in *Glast-Cre^ERT2^; Sun1:GFP* mouse retinal sections at 1 week, 3 weeks, and 4 months post-CCA injection.

**Supplementary Figure** **9. Analysis of p27^kip1^ expression at different days post-CCA injection on P6**

(A) Experimental design. (B-D) Analysis of EdU incorporation co-labeled with p27^kip1^ and MG-specific marker Sox2 in the retinas received two EdU injections on D1-2 (C) and D11-12 (D) post-CCA treatment. Control retinas received EdU injections on D11-12 without CCA treatment on D0.

**Supplementary Figure** **10. Analysis of p27 expression at different days post-CCA injection on P28.**

(A) Experimental design. (B-E) Immunohistochemical analysis of p27^kip1^ expression in *Glast-Cre^ERT2^; Sun1:GFP* mouse retinal sections at 1 week, 3 weeks, and 4 months post-CCA injection.

**Supplementary Figure** **11. Preprocessing and filtering of scRNA data and removal of doublet cells**

(A) UMAP plot showing the clustering of control, CCA, and CCANT-treated cells. (B) Feature plot showing the expression of tdTomato. (C) UMAP plot and clustering of control, CCA, and CCANT-treated cells after removal of tdTomato-negative cells. (D) Feature plot showing tdTomato expression after tdTomato-negative cell removal. (E) UMAP plots of control, CCA, and CCANT groups separately. (F) Proportions of control, CCA, and CCANT groups in the UMAP cell clusters shown in (C). (G) Feature plot indicating doublets in pink. (H) Proportion of doublets in all cell clusters, with cluster 9 showing over 70% doublets, indicating low quality. (I) Clustering of control, CCA, and CCANT-treated cells with cluster labels C4-C9. (J) Proportion of control, CCA, and CCANT groups in cell clusters after doublets removal.

**Supplementary Figure** **12. Removal of possibly contaminated cells**

(A) UMAP plot showing seven cell clusters after removing doublet cells and small clusters composed by non-MG cells (C4-9 in Supplementary Figure 10I,J). (B) Percentages and numbers of cells in three groups across seven clusters. Two clusters of MG (C3 and C4) express rod genes such as *Rho*, *Gnat1*, and *Nrl*. In the C4 cluster, the percentages of cells from each group are similar, suggesting that C4 is likely a cluster of MG cells contaminated with rod processes during retinal cell isolation. This cluster was removed from further analysis. The C3 cluster was enriched in CCA or CCANT-treated samples and was kept for further analysis. (C) Split feature plots showing that the cells in the cluster C4 exhibited similar proportional contributions from all three groups and expressed lower level of rod-specific genes such as *Rho* than C3*.* C4 was defined as contaminated cells and removed for further analysis. (D) UMAP plot showing the cells in the G1, G2/M or S phase. (E) Proportions of cells in different cell cycle phases across six clusters.

**Supplementary Figure** **13. Characterization of the rod-MG and rod clusters in the scRNA-seq analysis**

(A) UMAP plots showing expression patterns of various rod genes.

**Supplementary Figure** **14. RNA *in situ* hybridization analysis of rod gene expression in dissociated MG following CCA treatment**

(A) Schematic of RNA *in situ* hybridization analysis for rod gene expression in dissociated MG 3 weeks post-CCA treatment. (B-C) Expression levels of rod mRNAs (*Rho and Gnat1*) in dissociated retinas from control and CCA-treated groups. (B1-C2) Enlarged views of boxed regions in (B-C). (D) Quantification of *Gnat1* and *Rho* mRNA expression by dots per Sun1:GFP+ cell. Data presented as mean ± SEM,total 300 GFP+ cells were measured in two biological sample per group. *****P* < 0.0001, (unpaired two-tailed student’s t-test).

**Supplementary Figure** **15. Expression level changes of MG genes in reactivated MG**

(A) Feature plots showing the expression patterns of genes associated with the quiescent state (*Kcnj10, Glul, Rlbp1,* and *Aqp4*) and the reactivated state of MG (*Vim* and *GFAP*). (B) Box plots showing expression levels of MG genes in different clusters. (C) Box plots showing expression levels of MG genes in different groups.

**Supplementary Figure** **16. Absence of neurogenic progenitor clusters induced by CCA treatment**

(A) Feature plots illustrating neurogenic or progenitor genes, Notch pathway genes, *Ccnd1*, *Cdkn1b*, and *Btg2*.

**Supplementary Figure** **17. Detection of MG-derived HuC/D+ cells in the INL after CCA treatment**

(A) Experimental design. (B-C) Representative images of control uninjected retinas or retinas infected by CCA virus. EdU was co-labeled with amacrine cell marker HuC/D. (D-E) Enlarged views of the boxed area in (B,C). (F) Quantification of tdT+ EdU+ HuC/D+ cells as a percentage of total tdT+ EdU+ cells at different time points post-CCA treatment in *Glast-Cre^ERT2^; tdTomato* retinas. Data presented as mean ± SEM, n=3. *****P* < 0.0001 (one-way ANOVA with Tukey post hoc test).

**Supplementary Figure** **18. Absence of TUNEL-positive MG in the retina following CCA treatment**

(A) Experimental design. (B-C) Representative images of TUNEL analysis of *Glast-Cre^ERT2^; Sun1:GFP* retinas in control (B) or CCA-treated retinas (C) at three weeks after CCA treatment.
